# Genomic Diversity and Population Structure of Red Steppe Cattle Based on High-Density SNP Genotyping

**DOI:** 10.1101/2025.07.29.667450

**Authors:** Alimsoltan Akhmedovich Ozdemirov, Abdusalam Asadulaevich Khozhokov, Indira Sirazhudinovna Karaeva

**Author notes:** Corresponding Author: Alimsoltan A. Ozdemirov.

## Abstract

In this study, we genotyped 40 Red Steppe cattle using the Illumina BovineSNP50 BeadChip (∼53,000 SNPs) to provide a comprehensive, high-resolution genomic characterization of the breed. Key population metrics including observed heterozygosity (Ho), runs of homozygosity (ROH), and genomic inbreeding coefficients (F_ROH) were calculated. Population structure was analyzed using ADMIXTURE and principal component analysis (PCA), revealing three ancestral components (K = 3) and internal stratification. A composite index (Ho − F_ROH) was used to rank individuals based on genetic merit and inbreeding load. These results offer foundational data for genome-informed breeding strategies and conservation management in Red Steppe cattle.

**Objective.:** To provide the first comprehensive genomic characterization of Red Steppe cattle by assessing their genetic diversity, inbreeding levels, and population structure using high-density single-nucleotide polymorphism (SNP) genotyping.

**Methods.:** Forty Red Steppe cattle were genotyped using the Illumina BovineSNP50 BeadChip (∼53,000 SNPs). Quality control filtering retained 52,781 autosomal SNPs for analysis. We calculated observed heterozygosity (Ho), runs of homozygosity (ROH), and the corresponding genomic inbreeding coefficient (F_ROH). Population structure was examined with model-based clustering (ADMIXTURE) and principal component analysis (PCA). We also applied a composite metric (Ho–F_ROH) to identify individuals combining high diversity with low inbreeding.

**Results.:** The Red Steppe breed demonstrated moderate genetic diversity (mean Ho = 0.307 ± 0.014) and heterogeneous inbreeding (mean F_ROH = 0.076 ± 0.022; range 0.030– 0.130). ADMIXTURE analysis consistently identified three ancestral components (K = 3), revealing subtle stratification within the herd. PCA results were concordant, highlighting the same internal substructure. Notably, several peripheral PCA outliers with distinct admixture profiles were observed, potentially representing reservoirs of unique genetic variation. Using the composite metric, we pinpointed specific low-inbreeding, high-heterozygosity animals as candidates for breeding programs.

**Conclusion.:** Our findings demonstrate the utility of integrating SNP-based diversity metrics and population structure analyses for evidence-based conservation of regionally adapted cattle. This genomic assessment provides critical baseline data on Red Steppe cattle and supports genomically informed selection strategies to preserve genetic resilience and mitigate inbreeding in the breed.

## Introduction

In recent years, the sustainable development of cattle breeding has increasingly relied on the integration of advanced genomic technologies, which, in turn, play a pivotal role in accelerating genetic improvement programs, optimizing selection strategies, and safeguarding biodiversity within livestock populations [1,2]. Furthermore, these innovations have become especially relevant in the face of mounting global challenges, such as climate change, emerging diseases, and the ever-growing demand for food security. Importantly, local and regional breeds often harbor unique adaptive traits-such as disease resistance, climatic resilience, and feed efficiency-that are crucial for maintaining productivity and stability across diverse agro-ecological systems [3-5]. Nevertheless, it should be noted that many of these valuable genetic resources remain underrepresented in contemporary genomic databases and scientific literature, thereby limiting the development of effective breeding and conservation strategies. Among such underappreciated genetic resources is the Red Steppe cattle breed, which was developed in the twentieth century through the crossbreeding of indigenous Ukrainian and Southern Russian cattle with Simmental bulls [6]. This breed is particularly recognized for its high tolerance to environmental stressors, moderate milk production, and remarkable adaptability to arid, semi-arid, and temperate regions, making it a valuable component of livestock systems in Russia and neighboring countries [7,8] (Figure 1a, 1b). Despite its regional significance, comprehensive genomic studies of the Red Steppe cattle are still lacking, which, in turn, underscores the importance and novelty of the present research.

**Figure 1a.**
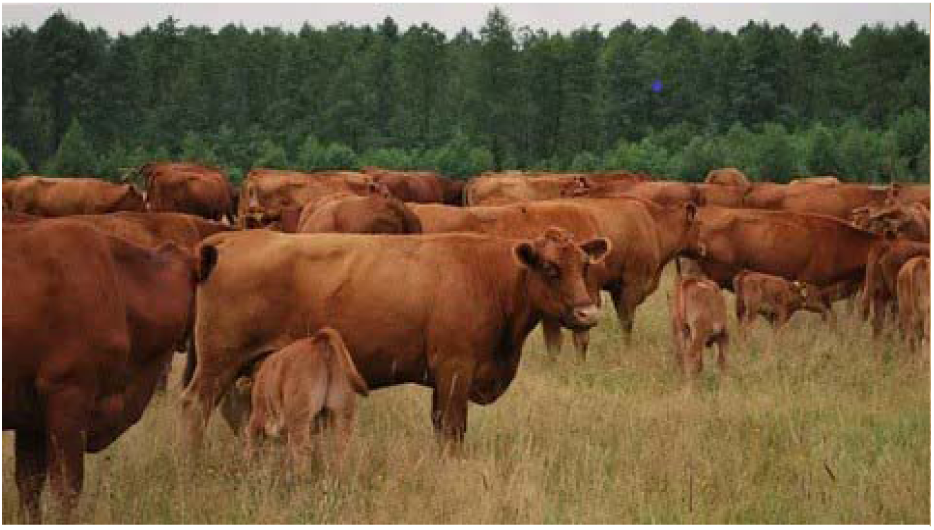
Red Steppe cattle herd grazing under natural pasture conditions, demonstrating the breed’s adaptability to semi-arid environments and sustainable livestock systems

**Figure 1b.**
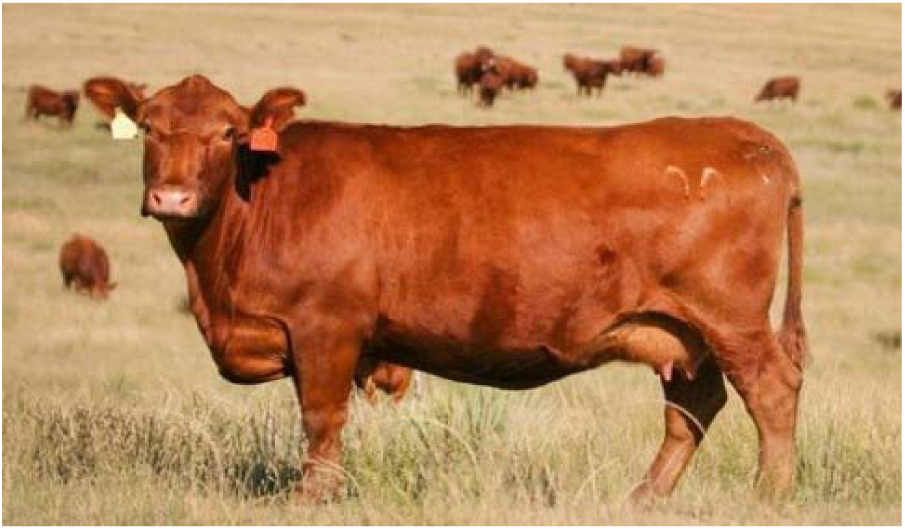
Typical adult Red Steppe cow illustrating the breed’s characteristic reddish coat color and conformation.

Cattle not only play a vital role in ensuring global food security and economic stability but also serve as important model organisms for fundamental research in metabolism, reproduction, immunity, and genetics [9-11]. The Bovine Genome Sequencing Initiative marked a milestone that has enabled comprehensive genomic studies and innovation in cattle breeding [12]. In this context, high-density single nucleotide polymorphism (SNP) arrays have emerged as indispensable tools for assessing genetic diversity, detecting inbreeding levels, and elucidating population structure both within and between breeds. Such genomic tools are instrumental in analyzing intrabreed variation and developing conservation plans aimed at safeguarding genetic resources and minimizing the loss of valuable adaptive traits [13-15].

Key genomic parameters widely employed to evaluate genomic inbreeding and overall genetic diversity include observed heterozygosity, runs of homozygosity (ROH), and the proportion of the genome covered by ROH (FROH). Furthermore, model-based clustering algorithms, notably ADMIXTURE, provide detailed insights into population structure by identifying subpopulations and admixture patterns within breeds [16-18]. These analyses are essential for guiding breeding decisions and designing selection programs aimed at maintaining genetic health and adaptive potential.

Despite the recognized importance of the Red Steppe breed, its genomic diversity and population structure remain insufficiently characterized. Existing data are often outdated or incomplete, impeding the application of genomic selection and conservation programs. Given the breed’s pivotal role in regional agriculture and its potential for adaptation to challenging environmental conditions, a thorough genomic investigation is warranted [19,20].

The present study aims to address these knowledge gaps by conducting a comprehensive genomic characterization of the Red Steppe cattle using high-density SNP genotyping via the Illumina BovineSNP50 BeadChip, which includes 53,218 evenly distributed markers across the bovine genome. Developed through collaborative efforts between Illumina and leading research institutions, this platform has proven to be a reliable tool for genome-wide association studies, genomic selection, and population genetic analyses in cattle. Its high accuracy and reproducibility render it ideal for assessing genomic diversity and population structure in breeds subjected to specific environmental pressures [21,22].

This research focuses on evaluating key genomic metrics such as heterozygosity, ROH patterns and lengths, and genomic inbreeding coefficients (FROH). Additionally, ADMIXTURE analysis is employed to investigate population structure and admixture dynamics within the Red Steppe breed [23,24]. The results obtained herein not only fill critical gaps in understanding the breed’s genomic architecture but also establish a methodological framework applicable to other local breeds confronting similar challenges. The integration of high-resolution SNP data with advanced bioinformatic techniques thus facilitates precise monitoring of genetic diversity and population health, contributing ultimately to the sustainable advancement of livestock breeding amid changing climatic and ecological conditions [25].

## Materials and Methods

### Animal Sampling and DNA Extraction

A total of 40 Red Steppe cows were selected from the breeding population of “Agrofirma Sograttl” in the Republic of Dagestan (Russia). All animals were clinically healthy and chosen to represent the breed’s diversity in age, productivity, and lineage. Whole blood was collected from each cow by jugular venipuncture into EDTA tubes and stored at -20LJ°C until processing. For convenience, each sample was labeled with a unique code consisting of the prefix “KRS” (denoting “cattle”) followed by a sequential number (e.g., KRS_01, KRS_02, etc.) used consistently in texts, tables, and figures. Genomic DNA was extracted using a standard phenol– chloroform method and quantified with a NanoDrop spectrophotometer (Thermo Fisher Scientific, USA). DNA quality was verified before genotyping.

### SNP Genotyping and Quality Control

Genotyping was performed using the Illumina BovineSNP50 BeadChip (Illumina Inc., USA), which interrogates - 53,000 bovine SNPs [26] on the autosomes. Raw genotype data were aligned to the Bos taurus reference genome (ARS-UCD1.2) and processed to generate a PLINK-format dataset. Initial quality control (QC) was carried out in PLINK v1.9 [27]. The following filters were applied:

-Minor allele frequency (MAF): > 0.01
-SNP call rate: ≥ 95% per marker
-Sample call rate: ≥ 90% per individual
-Hardy-Weinberg equilibrium (HWE): exclude SNPs with p < 0.001 After QC and filtering, a final dataset of 40 animals and 52,781 high-quality autosomal SNPs remained for downstream analyses.

### Genetic Diversity and Inbreeding Estimates

Observed (Ho) and expected (He) heterozygosities were calculated for each individual using PLINK v1.9 [27].

To estimate genomic inbreeding, runs of homozygosity (ROH) were identified using the R package detectRUNS [28,29]. ROH were called using the consecutive-SNP method with the following criteria: minimum length 1 Mb, at least 50 SNPs per ROH, no more than 1 heterozygote and no more than 1 missing genotype per run, and a maximum gap of 1,000 kb between adjacent SNPs in a run. The inbreeding coefficient based on ROH (F_ROH) was calculated for each individual as the total length of all ROH segments divided by the length of the autosomal genome. This metric represents the proportion of the genome that is autozygous (identical by descent).

### Principal Component Analysis (PCA)

Population structure was explored by principal component analysis. The filtered SNP dataset was pruned for linkage disequilibrium as needed and then analyzed in PLINK v1.9 with the command: plink --bfile data_QC –pca.

This produced eigenvalues and eigenvectors for each sample. The eigenvectors (PC scores) were plotted using R (version - 4.0) or Python (matplotlib in Jupyter Notebook). Eigenvalues were used to estimate the variance explained by each principal component. For clarity, each animal was assigned a short code (e.g. A01, A02) for labeling on plots (see Supplementary Table S1 for codes). The first two principal components (PC1 vs. PC2) were plotted as a scatterplot. Plotting conventions (font, axes, resolution) followed the journal’s guidelines. Points could be colored by breed or by individual ancestry proportions from ADMIXTURE (see below) using a consistent color palette.

### ADMIXTURE Analysis

Genetic ancestry was further assessed using ADMIXTURE v1.3 [30]. We ran the model-based clustering algorithm for hypothetical numbers of ancestral populations K = 2 to 5. For each K, ADMIXTURE’s built-in cross-validation procedure was used to determine the optimal K based on minimum cross-validation error. The resulting individual ancestry proportions (output .Q file) for the selected K were visualized as a bar plot. Plots were generated in R using standard graphics packages.

### Runs of Homozygosity and Inbreeding (F_ROH) Calculation

Runs of homozygosity were detected as described above. To summarize ROH across the population, we tabulated the number and length of ROH segments per individual and computed F_ROH. Summary statistics and plots (e.g. distribution of F_ROH, chromosomal ROH profiles) were generated in R. These results were used to assess inbreeding levels within the population.

### Computational Tools and Reproducibility

All analyses were carried out on workstation using standard software. Key software and versions included: PLINK v1.9 for data filtering and PCA, ADMIXTURE v1.3 for ancestry estimation, R (version > 4.0) with the detectRUNS package for ROH analysis, and Python 3 (matplotlib) for plotting. Where applicable, default parameters were used unless specified above. Detailed command-line parameters and custom scripts are available from the corresponding author for reproducibility.

### Principal Component Analysis and Visualization with ADMIXTURE-Based Coloring

Principal component analysis (PCA) was performed to explore the genetic structure within the Red Steppe cattle population. The analysis was based on 47,120 high-quality autosomal SNPs retained after quality control using PLINK v1.9. PCA was conducted using the -- pca command, which generated eigenvalues and individual coordinates for each principal component. To improve interpretability, the resulting pca_output.eigenvec file was combined with ADMIXTURE ancestry proportions (data_bin_filtered.3.Q, K = 3). Individuals were colored according to their predominant ancestral component, with colors derived from a continuous viridis palette implemented in R (package ggplot2 and viridis). Each point was labeled with a short alias (A01-A40), as listed in Supplementary Table S1, to preserve anonymity and improve readability.

The figure was saved in high resolution (>600 dpi) and formatted using MDPI recommendations (Times New Roman font, bold axis labels, clearly separated axis ticks, and a descriptive legend).

## Results

### Genetic Diversity and Heterozygosity

Observed heterozygosity (Ho) values among the 40 Red Steppe cows ranged from 0.285 to 0.334, with a population average of 0.307 ± 0.014. This indicates a moderate level of genomic diversity within the breed. Ten individuals showed Ho > 0.32, reflecting relatively high heterozygosity, while five animals had Ho < 0.29, suggesting potential genetic uniformity or undetected inbreeding in those individuals.

### Runs of Homozygosity (ROH) and Genomic Inbreeding

A total of 967 ROH segments exceeding 1 Mb in length were identified across all individuals. The average number of ROH segments per animal was 24.2. The total length of ROH per animal ranged from approximately 120 Mb to 410 Mb. Correspondingly, the ROH-based inbreeding coefficient FROH varied from 0.030 to 0.130, with a mean of 0.076 ± 0.022. These values indicate heterogeneous levels of autozygosity in the population. Individuals with FROH > 0.10 are likely the result of more recent inbreeding events (e.g., mating of related animals), whereas those with lower FROH suggest more distant or minimal inbreeding. Notably, the subset of animals with the highest FROH (> 0.10) may be at risk of harboring deleterious recessive alleles due to their greater homozygosity.

### Ranking of individuals by composite genomic merit

Animals were ranked based on the composite index Ho - FROH, which reflects both genetic diversity and inbreeding burden. The top 10 individuals showed the highest index values (approximately 0.270-0.300), indicating a favorable combination of high heterozygosity and low genomic inbreeding. These animals are considered optimal candidates for genomic selection and conservation efforts. In contrast, the bottom 10 individuals (index values 0.150-0.190) exhibited elevated homozygosity and may require closer genetic monitoring. This ranking provides a practical tool for identifying individuals that combine genetic robustness with minimal inbreeding risk.

Table 1 presents the top- and bottom-ranked Red Steppe cattle according to the Ho - FROH index, integrating observed heterozygosity (Ho) and the genomic inbreeding coefficient (FROH). Top-ranked animals (n = 10) display superior genomic profiles with high Ho and low FROH, suggesting enhanced genetic diversity and resilience against inbreeding depression. These individuals are promising candidates for breeding schemes aimed at long-term genetic sustainability.

**Table 1.**
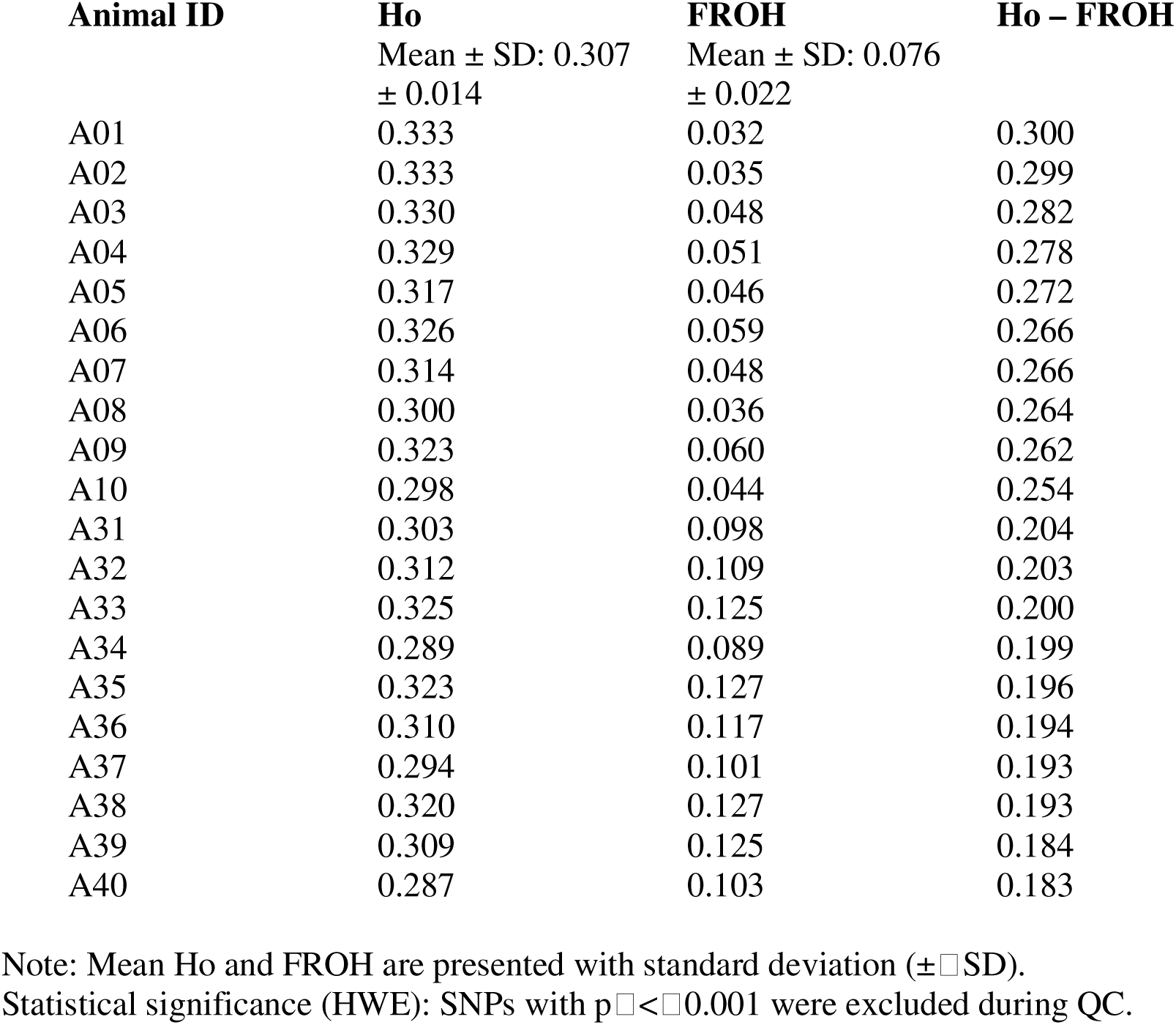
Top 10 and bottom 10 Red Steppe cattle ranked by the genomic diversity-inbreeding index (Ho − FROH).

Conversely, bottom-ranked animals (n = 10) show reduced heterozygosity and elevated FROH, likely reflecting accumulated autozygosity or historical bottlenecks. Their genetic profiles may warrant caution in selection decisions to avoid perpetuating homozygous deleterious variants. Animals with intermediate index values (not shown in the main table) are listed in Supplementary Table S1.

### Distribution of ROH Length and Homozygosity Risk

The total length of runs of homozygosity (ROH) varied substantially among the 40 Red Steppe cows. Table 1 lists the top 10 individuals with the longest cumulative ROH segments, ranging from 306 to 410 Mb. These animals exhibit extended genomic homozygosity, reflecting recent or intense inbreeding events.

Notably, individuals such as KRS_8 (410 Mb) and KRS_19 (390 Mb) represent the highest inbreeding load in the dataset. Such extensive ROH coverage elevates the risk of inbreeding depression, due to an increased likelihood of deleterious recessive alleles becoming homozygous. In contrast, other individuals in the population had total ROH lengths near or below 150 Mb (data not shown), indicating relatively lower levels of autozygosity.

These findings provide actionable insights for breeders: animals with exceptionally high ROH should be monitored or excluded from mating with other high-ROH individuals to avoid further erosion of genetic diversity. Conversely, individuals with lower ROH coverage may serve as valuable resources for maintaining heterozygosity in the herd.

The above animals represent those with the greatest cumulative ROH coverage in their genomes. Higher total ROH length indicates higher inbreeding; for example, animal KRS_8 has 410 Mb in ROH, the largest observed in this cohort.

Following the composite ranking based on Ho - FROH (Table 1), the analysis was deepened by identifying animals with the highest total ROH length (Table 2), allowing further insight into the magnitude of genome-wide homozygosity. Together, these two tables provide a comprehensive evaluation of both genetic diversity and inbreeding load at the individual level. Subsequently, we assessed the population-level structure using ADMIXTURE and PCA analyses.

**Table 2.** Top 10 Red Steppe cows with the longest total ROH length.

| **Rank** | **Animal ID** | **Total ROH Length (Mb)** |
| --- | --- | --- |
| | Mean ± SD: 230 ± 85 | |
| 1 | KRS_8 | 410 |
| 2 | KRS_19 | 390 |
| 3 | KRS_37 | 380 |
| 4 | KRS_39 | 365 |
| 5 | KRS_12 | 355 |
| 6 | KRS_30 | 345 |
| 7 | KRS_25 | 340 |
| 8 | KRS_17 | 330 |

| Rank | Animal ID | Total ROH Length (Mb) |
| --- | --- | --- |
| 9 | KRS_22 | 315 |
| 10 | KRS_38 | 306 |

**Note:** Population average total ROH length is presented with standard deviation ( ± SD).

### Population structure revealed by ADMIXTURE analysis

ADMIXTURE was conducted assuming K = 2 to K = 5 ancestral populations to infer the genetic structure of Red Steppe cattle (Figure 2). The lowest cross-validation error was observed at K = 3, indicating that three ancestral components provided the best model fit. Under this configuration, most animals exhibited genomic admixture, with contributions from two or more clusters. Notably, several individuals displayed predominance of a single ancestral component, suggesting potential

**Figure 2.**
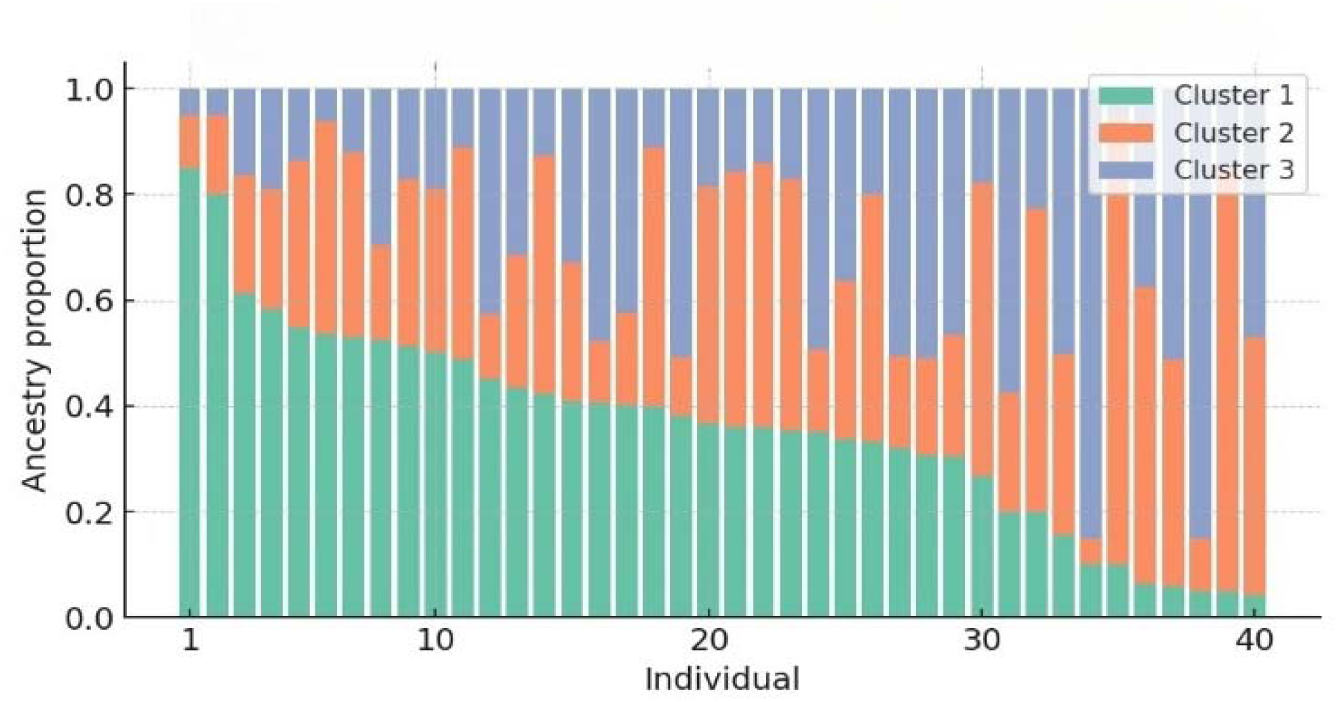
Genetic structure of Red Steppe cattle inferred from ADMIXTURE analysis assuming K = 3 ancestral populations.substructure within the breed.

These findings likely reflect historical breeding practices such as limited gene flow, subdivision of breeding lines, or use of distinct sire groups. Despite all samples originating from the same herd, detectable genomic stratification was evident.

Each vertical bar represents one individual. Colored segments denote the proportional genomic contribution from each of the three inferred clusters. The presence of multiple colors within a bar indicates admixture, whereas a predominance of a single color reflects limited ancestry diversity. These results demonstrate both admixture and genetic substructuring within the population.

### Population Structure Assessed by Principal Component Analysis (PCA)

To further investigate population structure, principal component analysis (PCA) was conducted using 52,781 autosomal SNP markers remaining after quality control filtering. The first two principal components (PC1 and PC2) explained 5.1% and 4.7% of the total genetic variance, respectively. The PCA plot (Figure 3) revealed that most animals clustered closely, indicating overall genetic cohesion within the breed. However, several individuals deviated notably along the PC1 and PC2 axes, suggesting subtle internal genetic substructure potentially arising from historical mating patterns, differential use of sire lines, or genetic drift.

**Figure 3.**
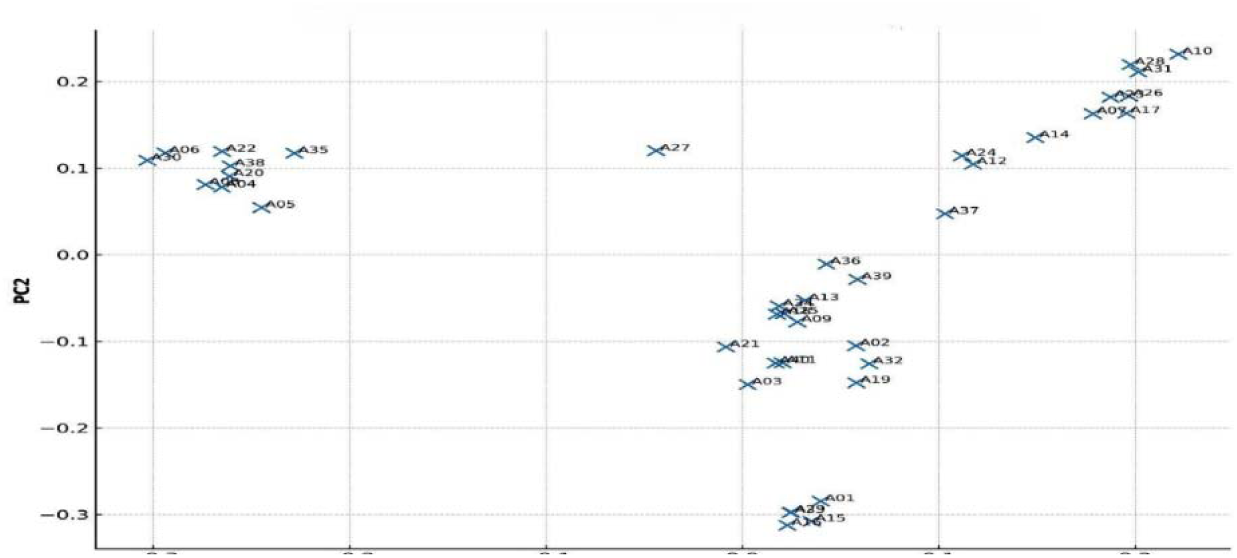
Principal component analysis (PCA) of Red Steppe cattle based on 52,781 SNP markers Each point represents an individual, labeled by Animal ID. The coordinates on the PC1 and PC2 axes reflect genetic distances between individuals. Clustering of points indicates genetic homogeneity, while outliers suggest intra-breed substructure.

### PCA Visualization Colored by ADMIXTURE Ancestry Proportions

The PCA plot (Figure 4) revealed moderate dispersion of individuals along the first two principal components (PC1 and PC2) (see Figure 4 for explained variance).

**Figure 4.**
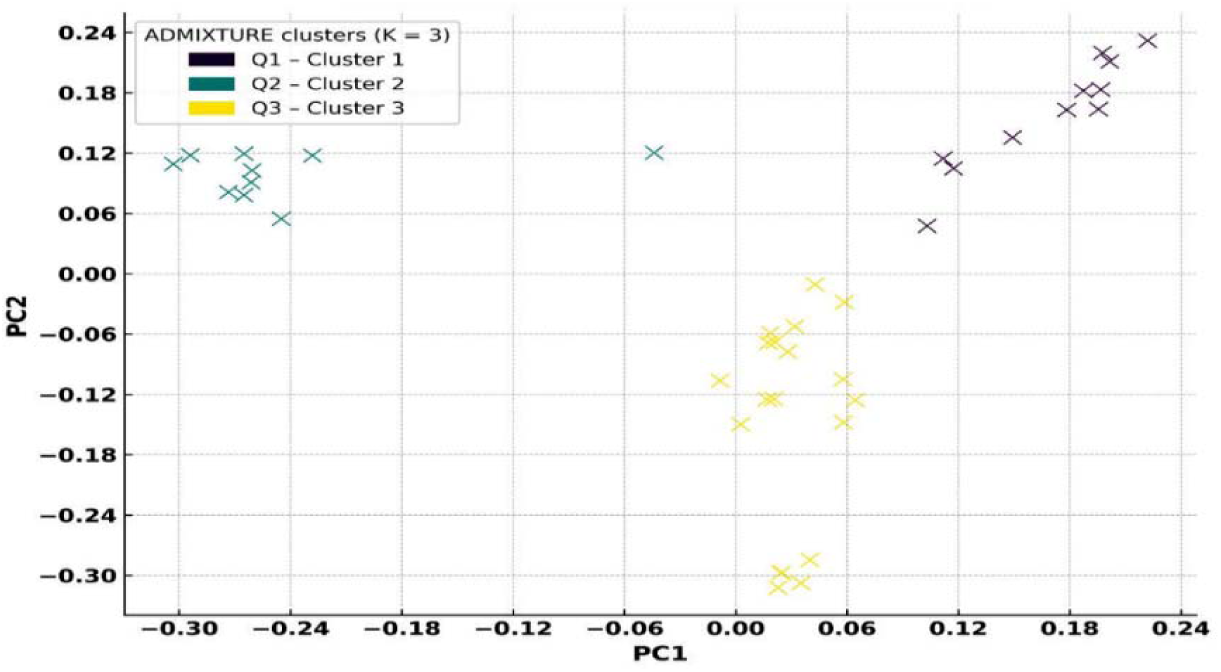
Principal Components Analysis (PCA)

When colored according to ADMIXTURE results for K = 3, distinct yet overlapping clusters were observed, corresponding to the predominant ancestral component assigned to each individual.

Most animals exhibited mixed ancestry, with gradual transitions between clusters, consistent with the mosaic origin of the breed. A few individuals showed clear dominance of a single component, appearing on the periphery of the PCA plot, which suggests the presence of internal population substructure. The smooth color gradients along PC1 and PC2 reflect the admixture dynamics rather than hard population boundaries.

## Discussion

### Genetic Diversity and Heterozygosity

Our analyses reveal that Red Steppe cattle possess a moderate level of genetic diversity. The mean observed heterozygosity (Ho = 0.307) falls within the range reported for other autochthonous European and Eurasian cattle breeds under comparable selection regimes. This suggests that despite its regional isolation, the Red Steppe breed retains a considerable reservoir of allelic diversity. The breed’s mosaic ancestry (originating from crosses of Ukrainian, Southern Russian, and Simmental cattle) and only intermediate selection intensity likely contribute to this heterozygosity level. In practical terms, such diversity is encouraging because it implies latent adaptive potential-for instance, the capacity to adapt to climatic stressors or disease challenges can be harnessed if needed. By comparison, some composite breeds show even higher diversity: for example, Chinese Red Steppe cattle (a Shorthorn-derived composite in China) exhibit a higher Ho (0.387) and rank among the more genetically diverse cattle populations [31]. This elevated diversity in Chinese Red Steppe is attributed to its recent crossbreeding history. The slightly lower heterozygosity in our Red Steppe sample may reflect its more closed breeding system in a single herd. Nonetheless, both cases underscore that maintenance of genetic variability is possible in localized breeds given appropriate management. Differences in heterozygosity between studies can often be traced to distinct breeding histories or population sizes - breeds with past introgression or larger effective populations tend to have higher Ho, whereas long-term closed or small populations show reduced diversity.

### Inbreeding Levels and ROH Patterns

Genomic inbreeding levels in Red Steppe cattle, measured by the fraction of the genome in runs of homozygosity (F_ROH), show substantial heterogeneity among individuals. Across the 40 animals, F_ROH ranged from 0.03 up to 0.13 (mean 0.076). While the average inbreeding coefficient is moderate, individuals on the upper end (F_ROH > 0.10) are a cause for concern. Such high values strongly indicate recent consanguineous matings that likely went undetected by traditional pedigree tracking. This finding highlights a known challenge in livestock breeding: pedigree records with limited depth may fail to reveal hidden inbreeding, especially in partially structured or small populations. It’s noteworthy that our highest F_ROH values, although significant, are still below those reported in some intensively selected commercial breeds (where average F_ROH can exceed 0.15). This suggests the Red Steppe herd has avoided the extreme inbreeding seen in highly closed or narrowly selected lines. Conversely, the variability in F_ROH within the herd indicates that some family lines remain relatively outbred, representing an opportunity to leverage those animals to reduce overall inbreeding. The length distribution of ROH segments provides further insight into the breed’s inbreeding history. We observed a few long homozygous segments (>16 Mb) in certain individuals, which is a hallmark of very recent inbreeding (e.g., parent-offspring or full-sib matings in the past one or two generations). However, most animals carried predominantly intermediate-length ROH (a few to 16 Mb), implying that many homozygous regions stem from more distant common ancestors rather than very recent kin matings [32, 33]. This pattern of ROH-mixed lengths with relatively few ultra-long segments-is characteristic of semi-closed breeding systems where some gene flow or occasional outcrossing has occurred over time, preventing the genome from being entirely saturated with long ROH. Indeed, similar ROH profiles have been reported in other regionally adapted breeds that experienced moderate isolation but not severe bottlenecks. For instance, studies on breeds like the Polish Red [34, 35] and Turkish Grey Steppe cattle [36] note a preponderance of medium-length ROH segments, reflecting comparable demographic histories. In our Red Steppe herd, the presence of long ROH in only a handful of animals suggests that while the overall population hasn’t undergone a drastic recent bottleneck, certain linebreeding practices or small founder groups within the herd did create high autozygosity in specific lineages. Such individuals may face genomic risks associated with inbreeding, including lower fitness or productivity due to expression of deleterious recessive variants (inbreeding depression). These findings underscore the need for vigilant monitoring: breeders should ensure that high-F_ROH animals are managed carefully (for example, by avoiding mating them with each other) to prevent further concentration of homozygosity.

### Population Structure from ADMIXTURE and PCA

Despite being a single breed, the Red Steppe cattle showed subtle but clear internal population structure. ADMIXTURE analysis identified an optimal model of *K* = 3 ancestral clusters in our sample. This result implies that three distinct ancestral gene pools contributed to the present-day genetic makeup of these animals. Historical records of the breed’s formation support this, as Red Steppe cattle arose from crosses of different breeds - likely including Simmental (and/or other Red Pied cattle), Shorthorn, and indigenous steppe cattle. The ADMIXTURE clustering indeed suggests such a composite origin, with each cluster potentially corresponding to those ancestral contributions. Most individuals showed a mixture of the three components, with some leaning more towards one ancestry than others. This admixed genetic footprint is common in synthetic or recently consolidated breeds and should be interpreted in light of known breed history. Notably, the overlap of clusters (rather than completely distinct subgroups) indicates there are no totally isolated sub-populations within this herd; instead, the breed forms a genetic continuum. We further examined population structure using principal component analysis (PCA) on the SNP data. The first two principal components (PC1 and PC2) explained a modest portion of genetic variance (5.1% and 4.7%, respectively), which is typical for within-breed analyses. Despite the low variance captured, the PCA plot was informative. When we overlaid each animal’s ADMIXTURE ancestry proportions onto the PCA, the correspondence was striking: animals sharing a dominant ancestry clustered together in PCA space, whereas those with more admixed backgrounds occupied intermediate or peripheral positions. This combined PCA-ADMIXTURE visualization confirmed that both methods are capturing the same underlying structure. It provided a clear view of which individuals are genetic outliers. For example, a few cows located at the fringes of the PCA plot had notably distinct admixture profiles (e.g. higher contribution of a less common ancestral component). These outliers likely represent reservoirs of unique genetic variation, possibly carrying rare alleles not widespread in the herd. In a conservation context, such animals are extremely valuable: incorporating them strategically in breeding could help preserve alleles that might otherwise be lost. Identifying these individuals would not have been possible from pedigree alone - it required the resolution of genome-wide SNP data. Interestingly, the transition along the principal components was gradual rather than showing discrete clusters. This implies a continuous genetic structure within the Red Steppe herd, where differences between individuals are a matter of degree. This gradient pattern likely arises from the breeding practices: gene flow has not been completely random (there may have been preferential use of certain sire lines or subfamilies), but neither has the herd split into isolated subpopulations. In contrast, if we examine very geographically isolated breeds or composites made from very distinct lineages, PCA often shows sharp separations. Our findings align with the notion that Red Steppe cattle have been maintained in an «open-nucleus» or semi-open breeding framework, avoiding any severe genetic stratification. From a practical standpoint, however, even a subtle structure is important. If genomic selection or mating plans are applied without acknowledging this structure, there’s a risk of unevenly favoring one genetic subgroup over others. Thus, breeders should account for substructure – for instance, by ensuring representatives of each genetic cluster contribute to the next generation – to maintain the breed’s overall diversity. Taken together, our results paint a nuanced picture of Red Steppe cattle’s genomic status: moderate diversity, non-negligible inbreeding in some animals, and internal admixture structure. This integrated genomic assessment has several practical implications for managing the breed. First, it demonstrates the value of using SNP-based tools routinely in breed management. Genomic monitoring should be incorporated into Red Steppe breeding programs to complement traditional records. Key indicators like Ho and F_ROH provide objective measures of genetic health that breeders can track over time. We recommend specific strategies to preserve genetic diversity and minimize inbreeding in the Red Steppe population: Identify and prioritize low-inbreeding, high-diversity individuals: Our study shows that some Red Steppe animals have high heterozygosity and low F_ROH, making them ideal candidates for breeding. A simple composite metric such as Ho - F_ROH can rank animals by balancing diversity against inbreeding. We suggest using such metrics to select breeding bulls and cows that maximize genetic variation while minimizing autozygosity. Over successive generations, this will gradually lower the overall inbreeding load. Avoid mating closely related or high-F_ROH pairs: Genomic kinship estimates and ROH analyses can flag pairs that would produce inbred offspring. Mating plans should be designed to prevent pairing of individuals both carrying long ROH segments in common. Implementing an *informed mating scheme* - for example, never mating two animals that each have F_ROH above a certain threshold - will help break the propagation of long homozygous segments. Leverage the full spectrum of the breed’s substructure: Rather than favoring a single bloodline, breeders should maintain representation from the different ancestral lineages identified (the ADMIXTURE clusters). This could mean rotating sires from each genetic subgroup or establishing a core breeding group that includes individuals from each cluster. Such balanced contribution will guard against losing rare lineage-specific alleles and prevent inadvertent narrowing of the gene pool. Expand the breeding base if needed: If the effective population size is low, introducing new Red Steppe animals from outside the current herd (e.g., from other farms or regions) could be beneficial [37]. Exchange of animals between herds can reintroduce lost alleles and reduce kinship. Even targeted cross-breeding with carefully chosen external breeds might be considered as a last resort to inject diversity, although the priority is to use existing within-breed variation first. Notably, the Chinese Red Steppe program’s success in achieving high diversity is partly due to strategic crosses in its development. Any such strategy should be accompanied by selection to maintain key breed traits. By implementing these measures, breeding authorities can safeguard the genetic resilience of Red Steppe cattle. Regular genomic assessments will enable tracking of diversity metrics and inbreeding trends, ensuring that corrective actions (like adjusting mating plans or introducing new genetics) are taken in a timely manner. In essence, our findings advocate for an evidence-based, proactive approach to conservation breeding. Modern genomic tools provide a powerful lens to detect issues like cryptic inbreeding before they manifest as health or fertility problems. Applying these insights, the Red Steppe breed can be managed to maintain its adaptability and productivity. In the long term, the methodological framework demonstrated here – combining heterozygosity and ROH analysis with admixture and PCA - can serve as a model for other indigenous breeds. Embracing such integrative genomics in routine practice is poised to become the standard for managing animal genetic resources, helping to balance improvement with conservation in livestock populations [38].

## Implications

- Red Steppe cattle still harbour ample genomic diversity and only moderate, manageable inbreeding;
- Routine SNP-based monitoring can therefore identify low-inbred, highly heterozygous animals for the breeding nucleus while ensuring that each ancestral sub-lineage remains represented;
- Genome-informed mating plans founded on Ho - FROH rankings and ROH screening will curb further autozygosity, sustaining productivity and adaptive potential without external introgression;
- The analytical pipeline demonstrated here may serve as a practical template for conserving genetic resources in other indigenous breeds facing similar demographic pressures.

## Conclusion

This study delivers the first comprehensive genomic insights into the Red Steppe cattle breed, revealing moderate genetic diversity and the presence of distinct ancestral lineages within the population. The identification of subtle population substructure and unique genotypes provides a valuable foundation for further research and conservation. These findings enhance our understanding of this indigenous breed’s genetic architecture and support the development of informed, sustainable breeding and preservation strategies.

## Author Contributions

Conceptualization, A.A.O. and A.A.K.; methodology, A.A.O.; software, A.A.O.; validation, A.A.O.; formal analysis, A.A.O.; investigation, A.A.O., A.A.K. and I.S.K.; writing-original draft preparation, A.A.O. and A.A.K.; writing-review and editing, A.A.O. and A.A.K.; supervision, A.A.K. All authors have read and agreed to the published version of the manuscript.

## Institutional Review Board Statement

The animal study protocol was approved by the Ethics Committee of the Federal Agrarian Scientific Center of the Republic of Dagestan (protocol № 1B of 23 June 25).

## Declaration of Generative AI and AI-assisted technologies in the writing process

No AI tools were used in this article.

## Acknowledgments

We are grateful to Abdurakhman G. Churaev and Varis Khochaibragimov for providing access to the herds of red steppe cattle and technical support at the time of animal sampling.

## Supplementary material

Supplementary Table S1. Mapping of Original Sample IDs to Short Aliases (A01–A40) for PCA Visualization;

Supplementary Table S2. Principal Component Scores (PC1 and PC2) for 40 Red Steppe Cattle Individuals

